# Changes in lung sialidases in male and female mice after bleomycin aspiration

**DOI:** 10.1101/2022.09.29.510121

**Authors:** Darrell Pilling, Kyle Sahlberg, Wensheng Chen, Richard H. Gomer

## Abstract

**Aim of the study:** Sialidases, also called neuraminidases, are enzymes that cleave terminal sialic acids from glycoconjugates. In humans and mice, lung fibrosis is associated with desialylation of glycoconjugates and upregulation of sialidases. There are four mammalian sialidases, and it is unclear when the four mammalian sialidases are elevated over the course of inflammatory and fibrotic responses, whether tissue resident and inflammatory cells express different sialidases, and if sialidases are differentially expressed in male and females.

**Materials and Methods:** To determine the time course of sialidase expression and the identity of sialidase expressing cells, we used the bleomycin model of pulmonary fibrosis in mice to examine levels of sialidases during inflammation (days 3 - 10) and fibrosis (days 10 - 21).

**Results:** Bleomycin aspiration increased sialidase NEU1 at days 14 and 21 in male mice and day 10 in female mice. NEU2 levels increased at day 7 in male and day 10 in female mice. NEU3 appears to have a biphasic response in male mice with increased levels at day 7 and then at days 14 and 21, whereas in female mice NEU3 levels increased over 21 days. In control mice, the sialidases were mainly expressed by EpCAM positive epithelial cells, but after bleomycin, epithelial cells, CD45 positive immune cells, and alveolar cells expressed NEU1, NEU2, and NEU3. Sialidase expression was higher in male compared to female mice. There was little expression of NEU4 in murine lung tissue.

**Conclusions:** These results suggest that sialidases are dynamically expressed following bleomycin, that sialidases are differentially expressed in male and females, and that of the four sialidases only NEU3 upregulation is associated with fibrosis in both male and female mice.

## Introduction

Many secreted and cell-surface mammalian proteins are glycosylated, and many of the glycosylated structures have sialic acids as the monosaccharide at the end of the polysaccharide chain.^1-3^ Sialidases, which are also called neuraminidases, remove the terminal sialic acid from these glycoconjugates.^4, 5^ There are four known mammalian sialidases, NEU1, NEU2, NEU3, and NEU4, and these sialidases have different subcellular localizations and substrate specificities.^4-6^ Of the four sialidases, only NEU3 is expressed on the extracellular membrane.^6-9^

The lungs from both mouse models of lung fibrosis and patients with idiopathic pulmonary fibrosis (IPF) have increased levels of sialidase activity in the bronchoalveolar lavage fluid (BALF), increased levels of NEU3 protein in the BALF, increased levels of NEU1, NEU2, and NEU3 in the lung tissue, and extensive desialylation of glycoconjugates.^10-14^ Mice with adenovirus-mediated gene delivery of NEU1 into the lungs, or aspiration of NEU3 into the lungs, have increased inflammation and fibrosis, whereas mice treated with general sialidase inhibitors, mice treated with NEU1 or NEU3 specific inhibitors, and *Neu3*^*-/-*^ knockout mice show attenuated inflammation and fibrosis after bleomycin challenge.^12-18^ These data indicate that NEU1 and/or NEU3 are involved in inflammation and/or fibrosis. However, the specific roles of NEU1 and NEU3, as well as the other sialidases, in mouse models of pulmonary fibrosis are unclear.^11-17, 19^

In the mouse bleomycin-induced pulmonary fibrosis model, there is an initial inflammation of the lungs lasting from 3 to 10 days, followed by the appearance of fibrosis in the lungs starting at approximately day 10.^20-24^ To elucidate possible functions of sialidases, we examined the correlation of sialidase levels and localization in the lungs with the appearance of inflammation and fibrosis in the mouse bleomycin model. We find that NEU1 levels peak at day 14 in male and day 10 in female mice, NEU2 levels peak at day 7 in male and day 10 in female mice, and NEU3 levels appear to have a biphasic response in male mice with peaks at day 7 and then at days 14 and 21, whereas in female mice NEU3 levels increased over 21 days. Following bleomycin aspiration, NEU1 expression did not closely correlate with any single specific inflammation or fibrosis marker in male mice, and qualitatively correlated with CD11b positive inflammatory macrophages in the post-BAL lung tissue in female mice, NEU2 levels were associated with some markers of inflammation, and NEU3 levels were associated with both inflammatory and fibrosis markers. NEU1, NEU2, and NEU3, were expressed by epithelial, endothelial, alveolar, and immune cells. We also find that NEU4 levels are low in lung tissue and NEU4 is sometimes upregulated and sometimes not upregulated following bleomycin.

## Materials and methods

### Mouse model of pulmonary inflammation and fibrosis

This study was carried out in strict accordance with the recommendations in the Guide for the Care and Use of Laboratory Animals of the National Institutes of Health. The protocol was approved by the Texas A&M University Animal Use and Care Committee (IACUC 2020-0272). All procedures were performed under 4% isoflurane in oxygen anesthesia, and all efforts were made to minimize suffering. Animals were housed with a 12-hour/12-hour light-dark cycle with free access to food and water, and all procedures were performed between 09:00 and noon. To induce inflammation and fibrosis, 7-8 week old 20-25 g male and female C57BL/6 mice (Jackson Laboratories, Bar Harbor, ME) were given an oropharyngeal aspiration of 3 U/kg (equivalent to 0.06 U/20g mouse) bleomycin (2246-10; BioVision Incorporated, Milpitas, CA) in 50 µl of 0.9% saline, or oropharyngeal saline alone as a control, as previously described.^25, 26^ All the mice were monitored twice daily to observe any sign of distress. At the indicated time points, mice were euthanized by CO_2_ inhalation, and bronchoalveolar lavage fluid (BALF) and lung tissue was obtained and analyzed as described previously.^26, 27^

### Staining of bronchoalveolar lavage fluid (BALF) cells

BALF cells were counted and processed to prepare cell spots as described previously.^26, 28^ After air drying for 48 hours at room temperature, some of the cell spots were fixed and immunochemistry was performed as described previously.^26, 29^, using anti-CD3 (NB600-1441, rabbit clone SP7, Novus Biologicals, Centennial, CO) to detect T-cells, anti-CD11b (101202, clone M1/70, BioLegend, San Diego, CA) to detect blood and inflammatory macrophages, anti-CD11c (M100-3, clone 223H7, MBL International, Woburn, MA) to detect alveolar macrophages and dendritic cells, anti-CD45 (147702, clone I3/2.3, BioLegend) for total leukocytes, anti-Ly6g (127602, clone 1A8, BioLegend) to detect neutrophils, with isotype-matched irrelevant rat (BioLegend) and rabbit (Novus Biologicals) antibodies as controls. All antibody concentrations and buffers were described previously.^26, 29^ Using a 40x objective, at least 150 cells from each stained BALF spot were examined and the percent positive cells was recorded.

### Lung histology

After collecting BALF, the lungs from the mice were harvested and inflated with Surgipath frozen section compound (#3801480, Leica, Buffalo Grove, IL), frozen on dry ice, and stored at - 80°C. 10 μm cryosections of lungs were placed on Superfrost Plus glass slides (VWR) and were air dried for 48 hours. Immunohistochemistry was done as previously described ^28, 30^ using anti-CD3, anti-CD11b, anti-CD11c, anti-CD45, and anti-Ly6g, antibodies with isotype-matched irrelevant rat and rabbit as controls. Positively stained cells were counted from randomly selected fields and presented as the number of positive cells per mm^2^, as described previously.^28, 30^ Lung sections were also stained with Sirius red to detect collagen and analyzed as previously described.^14, 30^ Other investigators and ourselves have shown that histological analysis of fibrosis and collagen content by sirius red staining correlates well with collagen content as measured by antibody staining and hyproxyproline and sirius red (Sircol) assays.^14, 16, 30-34^

Lung section slides were also stained with 1 µg/mL rabbit polyclonal anti-NEU1 (TA335236; Origene, Rockville, MD), anti-NEU2 (TA324482, Origene) or (24523-1-AP; Proteintech, Rosemont, IL), anti-NEU3 (TA590228, Origene) or (27879-1-AP; Proteintech), or anti-NEU4 (AP52856PU-N, Origene) antibodies as previously described.^28, 30^ The anti-NEU3 (TA590228, Origene) was used at 0.5 µg/mL in PBS-BSA/500 mM NaCl/0.1% NP-40 alternative (EMD Millipore, Billerica, MA) for 60 minutes at room temperature, as described previously.^13, 14^ Slides were then incubated with 1 µg/mL biotinylated donkey F(ab’)_2_ anti-rabbit (711-066-152, Jackson ImmunoResearch Laboratories, West Grove, PA) secondary antibodies for 30 minutes at room temperature. Antibodies were detected with ExtrAvidin alkaline phosphatase (Vector Laboratories, Burlingame, CA), staining was developed with the Vector Red Alkaline Phosphatase Kit (Vector Laboratories), and sections were counterstained with hematoxylin, as previously described.^26, 28, 35^ To quantify neuraminidase staining, as Vector Red Alkaline Phosphatase is fluorescent above 560 nm.^36-38^, section images were captured with a 4x lens on a Nikon ECLIPSE Ti2 microscope using a Texas Red filter cube with a 540-580 nm excitation filter, a 593-668 nm emission filter, and a 585 nm dichroic mirror (Nikon, Melville, NY). Fluorescence intensity image quantification was done as previously described.^14, 26, 30^ Briefly, the red fluorescent image and the total area of the tissue, from a corresponding DIC image (Fig. S1), were used to calculate the area of tissue stained as a percentage of the total area of the tissue, using ImageJ software version 1.53n.^39^

### Immunofluorescence staining

Immunofluorescence staining was done as previously described.^14, 35^ Briefly, 10 µm lung cryosections were incubated with anti-sialidase antibodies as described above, and either 5 µg/mL rat anti-mouse CD45 (BioLegend) or rat anti-epithelial cell adhesion molecule (EpCAM; clone G8.8, BioLegend). After washing, slides were then incubated with 1 µg/mL donkey F(ab’)_2_ anti-rabbit Rhodamine-Red-X (711-296-152, Jackson ImmunoResearch), and 1 µg/mL donkey F(ab’)_2_ anti-rat Alexa Fluor 647 (712-606-153, Jackson ImmunoResearch) for 30 minutes at room temperature. Washed slides were mounted with mounting media containing DAPI (H-1500, Vector Laboratories). Images were captured with a Nikon ECLIPSE Ti2 microscope.

### Statistical Analysis

Statistical analysis was performed using Prism 7.05 (GraphPad Software, La Jolla, CA). Statistical significance between two groups was determined by t test, or between multiple groups using ANOVA with Dunnett’s post-test, and significance was defined as p<0.05.

## Results

### Time course of inflammation following bleomycin aspiration

To determine how immune cell recruitment and fibrosis in the lungs correlates with the expression and localization of sialidases in the lungs after aspiration of bleomycin, mice were treated with saline or bleomycin at day 0 by oropharyngeal aspiration, and then analyzed at days 3, 7, 10, 14, and 21. As observed previously, ^40-43^ male mice that received bleomycin have a significant drop in body weight from days 4 through 7, whereas female mice had a significant, but less severe, weight loss from days 6 through 9 (Figure 1A). Compared to female mice, the rate of weight loss from day 0 to day 7 was significantly faster in male as assessed by linear regression analysis (p=0.035; F test = 4.5).

**Figure 1:**
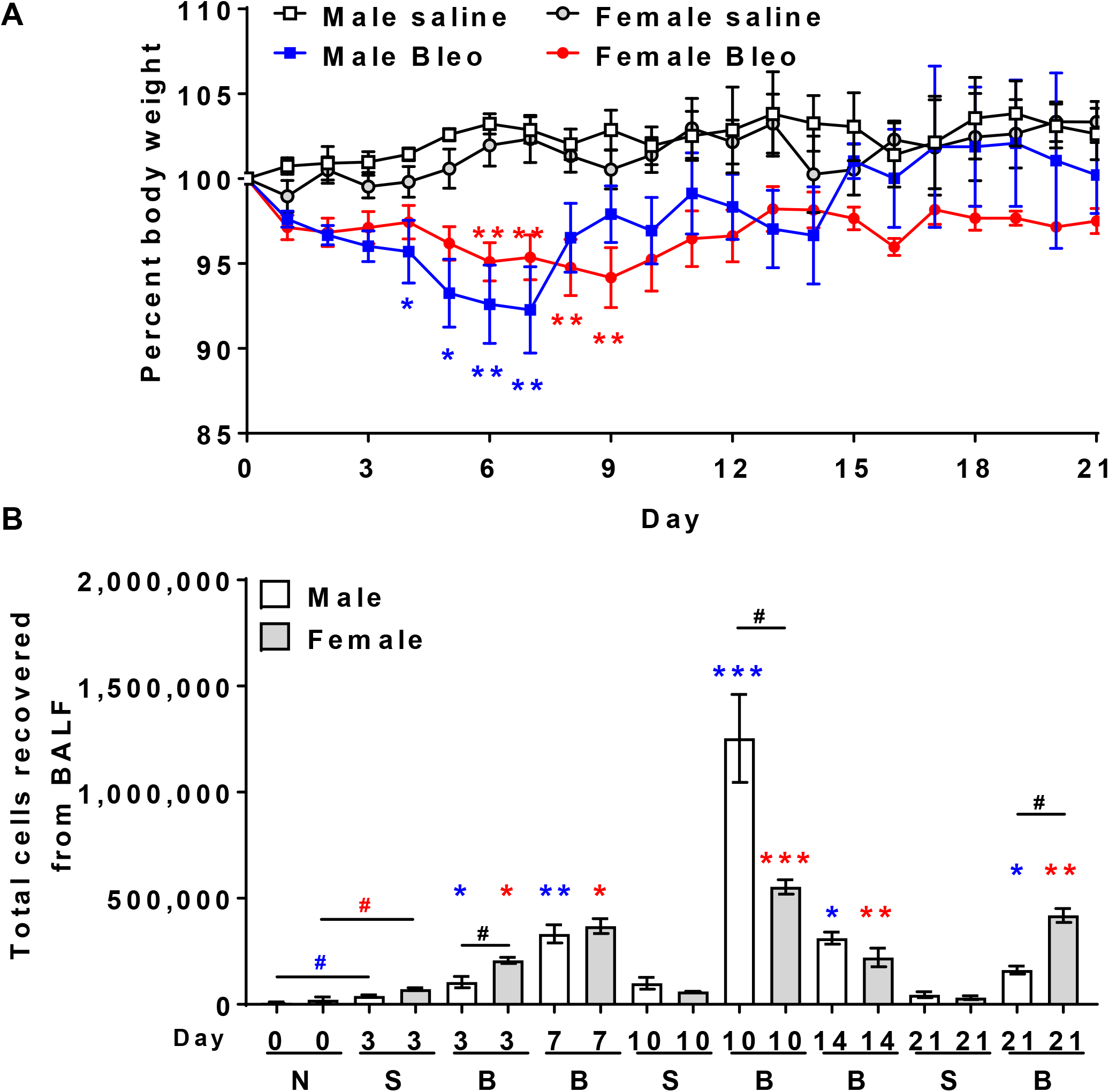
Changes in body weight and increased numbers of inflammatory immune cells in the BAL following bleomycin aspiration. **A)** Percent change in body weight in male and female mice after saline or bleomycin (Bleo) aspiration at day 0. Values are means ± SEM, n = 3-15 male or female mice per time point per group. *p < 0.05; **p < 0.01; (two-way ANOVA, Sidak’s multiple comparisons test). **B)** The total number of cells in male and female mouse BALF from naïve (N) untreated mice, and mice at days 3, 7, 10, 14, or 21, after saline (S) or bleomycin (B) aspiration. Values are means ± SEM, n = 3 male or female mice per group. *p < 0.05; **p < 0.01; ***p < 0.001 (one-way ANOVA, Dunnett’s test). In A and B, blue * or blue # indicate significance for male mice and red * or red # indicate significance for female mice, compared to day 3 saline treated male or female mice respectively. Black # p < 0.05 (t test between male and female mice at specific time points).

As previously observed,^14, 21, 26, 34, 42^ compared to saline controls, bleomycin aspiration led to an increase in the number of cells recovered from the BALF in both male and female mice over 21 days (Figure 1B). In the lungs of male and female mice that received bleomycin, compared to day 3 saline treated mice, there were increases in BAL cell counts at days 3, 7, 10, 14, and 21 (Figure 1B). Comparing bleomycin-treated male and female mice, there were more BALF cells in females at days 3 and 21, and more BALF cells in males at day 10 (Figure 1B). Compared to naïve mice, both male and female day 3 saline treated mice had more BALF cells (Figure 1B).

As previously observed, ^26, 34, 44^ bleomycin aspiration upregulated inflammatory cell counts in mouse lung BALF (Figure S2).^26, 34, 44^ In male mice, bleomycin increased the number of BALF CD3 T cells at day 10; CD11b inflammatory macrophages at days 7 and 10; CD11c resident alveolar macrophages at days 10, 14, and 21; Ly6g neutrophils at days 7 and 10; and CD45 positive immune cells from days 7 through 21, peaking at day 10 (Figures S2A-E). In female mice, bleomycin aspiration did not significantly increase BALF CD3 T cells, increased the number of BALF CD11b inflammatory macrophages at days 7 and 10; CD11c resident alveolar macrophages at days 7, 10, and 21; Ly6g neutrophils at day 7; and CD45 positive immune cells at days 7 through 21 (Figures S2A-E). Compared to naïve mice, both male and female day 3 saline treated mice had significantly more CD11c and CD45 positive BALF cells (Figures S2C and E).

After removing BALF, lung sections were stained to detect immune cells that were not removed by lavage. In male mice, bleomycin increased the number of lung tissue CD3 T cells at days 7, 10, and 14; CD11b inflammatory macrophages at days 7, 14, and 21; CD11c resident alveolar macrophages at days 7 through 21; and CD45 positive immune cells at days 7 through 21 (Figures S2F-J). In female mice, bleomycin did not significantly increase lung tissue CD3 T cells, increased lung tissue CD11b inflammatory macrophages at day 10; CD11c resident alveolar macrophages at days 7, 10 and 21; and CD45 positive immune cells at days 7 through 21 (Figures S2F-J). Bleomycin did not significantly affect Ly6g positive tissue neutrophils in either male or female mice. Compared to naïve mice, day 3 saline treated male mice had fewer CD3 and more Ly6g positive lung tissue cells (Figures S2F and I). Together, the results indicate that saline aspiration causes a slight amount of inflammation in the lungs, confirm that in male and female mice, bleomycin induces an inflammatory response, indicate that this response is greater in male than female mice, and allow a comparison of inflammation to the expression of sialidases described below.

### Time course of fibrosis following bleomycin aspiration

Lung sections were also stained with picrosirius red to detect total collagen. In male and female mice, aspiration of bleomycin, but not saline, increased picrosirius red staining (Figures 2A-K). Bleomycin aspiration increased picrosirius red staining in male mice at days 10, 14, and 21 (Fig. 2C-E and 2K), and in female mice at days 7 through 21 (Figures 2G-K). As previously observed, ^40, 42^ male mice treated with bleomycin had more fibrosis than female mice (Figure 2K).

**Figure 2.**
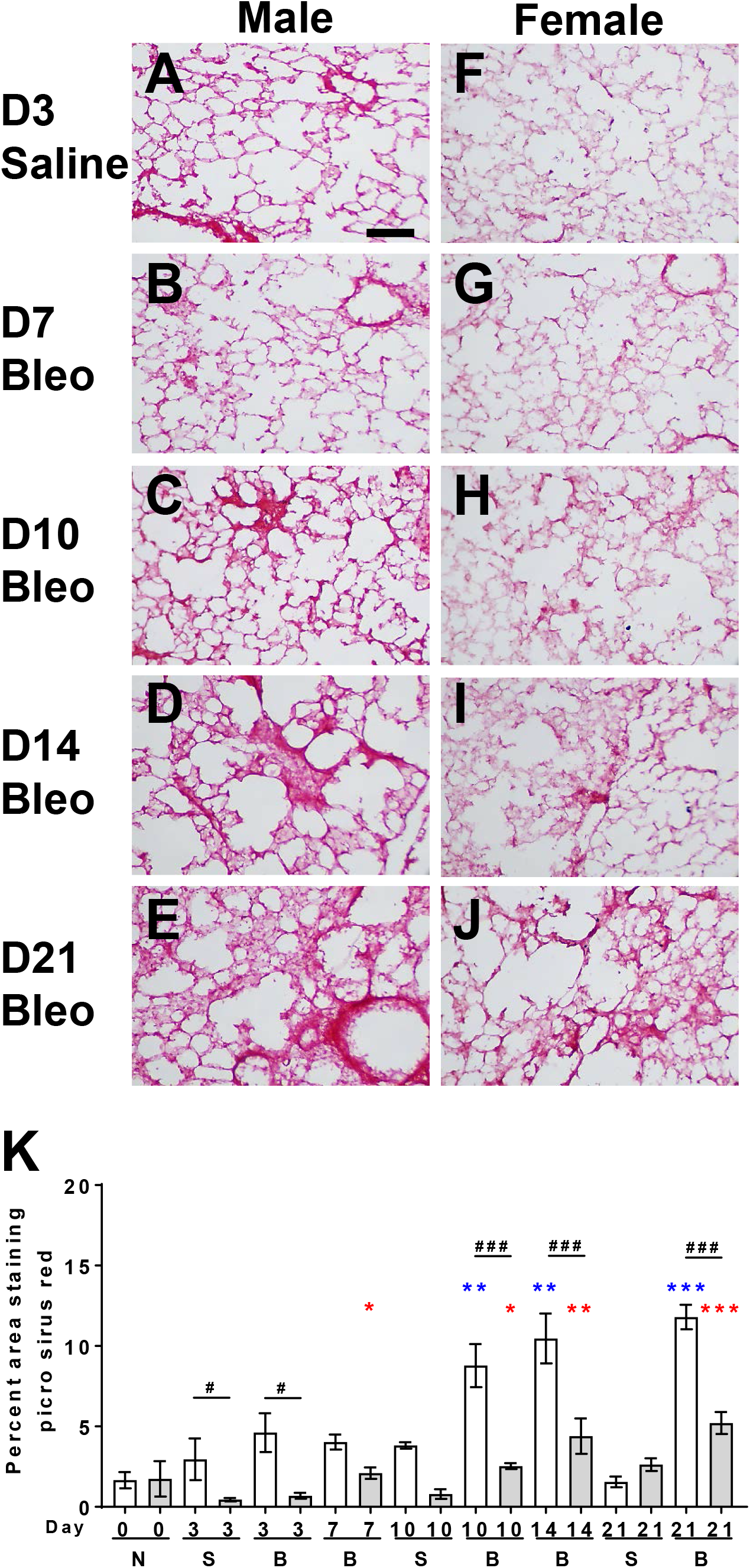
Induction of fibrosis in male and female mice following bleomycin. Sections of lung tissue were stained with picrosirius red to show collagen content. **A-E)** Male and **F-J)** female mice treated with saline at **A** and **F)** day 3, or bleomycin at **B** and **G)** day 7, **C** and **H)** day 10, **D** and **I)** day 14, or **E** and **J)** day 21. All images are representative of three mice per group. Bar is 0.1 mm. **K)** Three randomly chosen fields of view (each 0.275 mm^2^) were imaged for naïve (N) untreated mice, and mice at days 3, 7, 10, 14, or 21, after saline (S) or bleomycin (B). The percent area of the tissue showing staining was measured and the average was calculated. Values are mean ± SEM, n=3 male or 3 female mice. Blue asterisks indicate significance for male mice and red asterisks indicate significance for female mice, compared to day 3 saline treated mice. *p < 0.05; **p < 0.01, ***p < 0.001 (one-way ANOVA, Dunnett’s test). # p < 0.05, ### p < 0.001 (t test between male and female mice).

### NEU1 levels are high during the later stages of fibrosis in male mice, and are high only at day 10 in female mice

Compared to day 3 saline aspiration, bleomycin increased NEU1 staining of lung tissue at day 14 and 21 for male mice and day 10 for female mice (Figures 3A, S3 and S4). At all other time points, NEU1 staining in bleomycin-treated lungs was not significantly different from day 3 saline controls. In males, these NEU1 levels do not closely correlate with any of the inflammation or fibrosis markers (Figures 1B, S2, and 2). In females, the NEU1 levels correlate with CD11b positive inflammatory macrophages in the post-BAL lung tissue (Figure S2G). Compared to naïve male mice, day 3 saline treated male mice had significantly more NEU1 staining (Figures 3A and S5), indicating that saline aspiration can induce NEU1 expression in male mice. Together, the results suggest that increased NEU1 is associated with the later stages of fibrosis in the 21-day bleomycin model in male mice, but only some aspects of inflammation in female mice.

**Figure 3:**
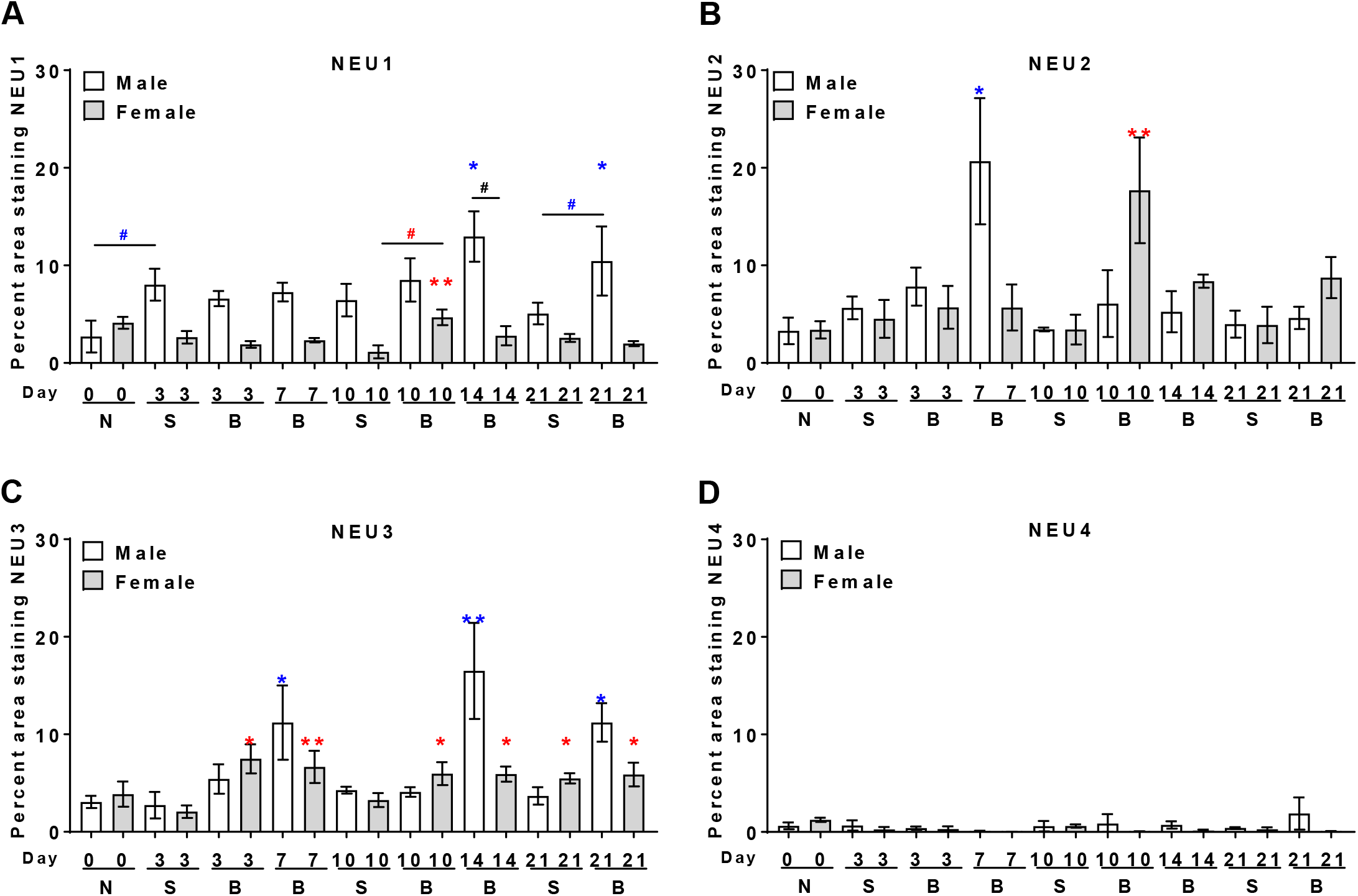
Changes in sialidase expression following bleomycin challenge. Cryosections of male and female mouse lungs were stained with anti-NEU1, NEU2, NEU3, or NEU4 antibodies. Three randomly chosen fields of view (each 2.25 mm^2^) were imaged, and the percent area of the tissue showing staining was measured and the average was calculated. Values are mean ± SEM, n=3 males and 3 females. *p < 0.05; **p < 0.01 (one-way ANOVA, Dunnett’s test). Blue asterisks indicate significance for male mice and red asterisks indicate significance for female mice compared to day 3 saline treated mice. # p < 0.05 (t test).

### NEU2 levels increase only during some stages of inflammation

Compared to day 3 saline controls, bleomycin aspiration led to a significant increase in NEU2 staining at day 7 in male mice, and day 10 in female mice (Figures 3B, S3, and S4). In males, the NEU2 levels only somewhat correlate with Ly6g positive BALF neutrophils (Figures 1B, S2D, and S3). In females, the NEU2 levels qualitatively correlate with NEU1 levels (Figure 3A) and CD11b positive inflammatory macrophages in the post-BAL lung tissue (Figure S2G). These results suggest that increased NEU2 levels are associated with inflammation but not fibrosis.

### NEU3 levels increase during inflammation and the later stages of fibrosis

Compared to day 3 saline controls, bleomycin aspiration led to a significant increase in NEU3 staining at days 7, 14, and 21 (but not day 10) in male mice, and days 3, 7, 10, 14, and 21 in female mice (Figures 3C, S3, and S4). In males, the increased NEU3 levels qualitatively correlate with CD11b positive inflammatory macrophages in the post-BAL lung tissue (Figure S2G). In females, the NEU3 levels qualitatively correlate with the total cells in the BALF (Figure 1B). In both male and female mice, increased NEU3 levels are also associated with the later stages of fibrosis in the 21-day bleomycin model.

### NEU4 levels did not increase in this model

We previously observed increased levels of NEU4 following bleomycin aspiration in female but not male mice.^13, 14^ However, in the current study, there was little NEU4 detected in the lungs of either male or female mice, and there was no significant increase following bleomycin (Figures 3D, S3, and S4). Together, these results suggest that unknown factors influence whether or not bleomycin aspiration causes an increase in NEU4 levels.

### Sialidase expression in epithelial, endothelial, and immune cells

NEU1 and NEU3 have previously been detected in in human and mouse lung epithelial cells by immunocytochemistry.^11, 12, 14, 45^ To determine the localization of sialidases at different times after bleomycin treatment, post-BAL lung sections were stained by immunofluorescence. Lung bronchiole epithelial vessels were identified by a columnar epithelium morphology and EpCAM staining. ^46^

As discussed above, NEU1 levels do not closely correlate with any of the inflammation or fibrosis markers. Although some NEU1 staining was localized to EpCAM positive cells (green arrows Figures 4B and 5B), along with EpCAM negative endothelial vessels (V’s in Figures 4 and 5), and occasional CD45 positive cells (* in Figures 4 and 5), most NEU1 positive cells within the lung alveoli did not localize with CD45, EpCAM, or EpCAM negative endothelial vessels (red arrows Figures 4 and 5, and unmarked in Figures 6-9).

**Figure 4:**
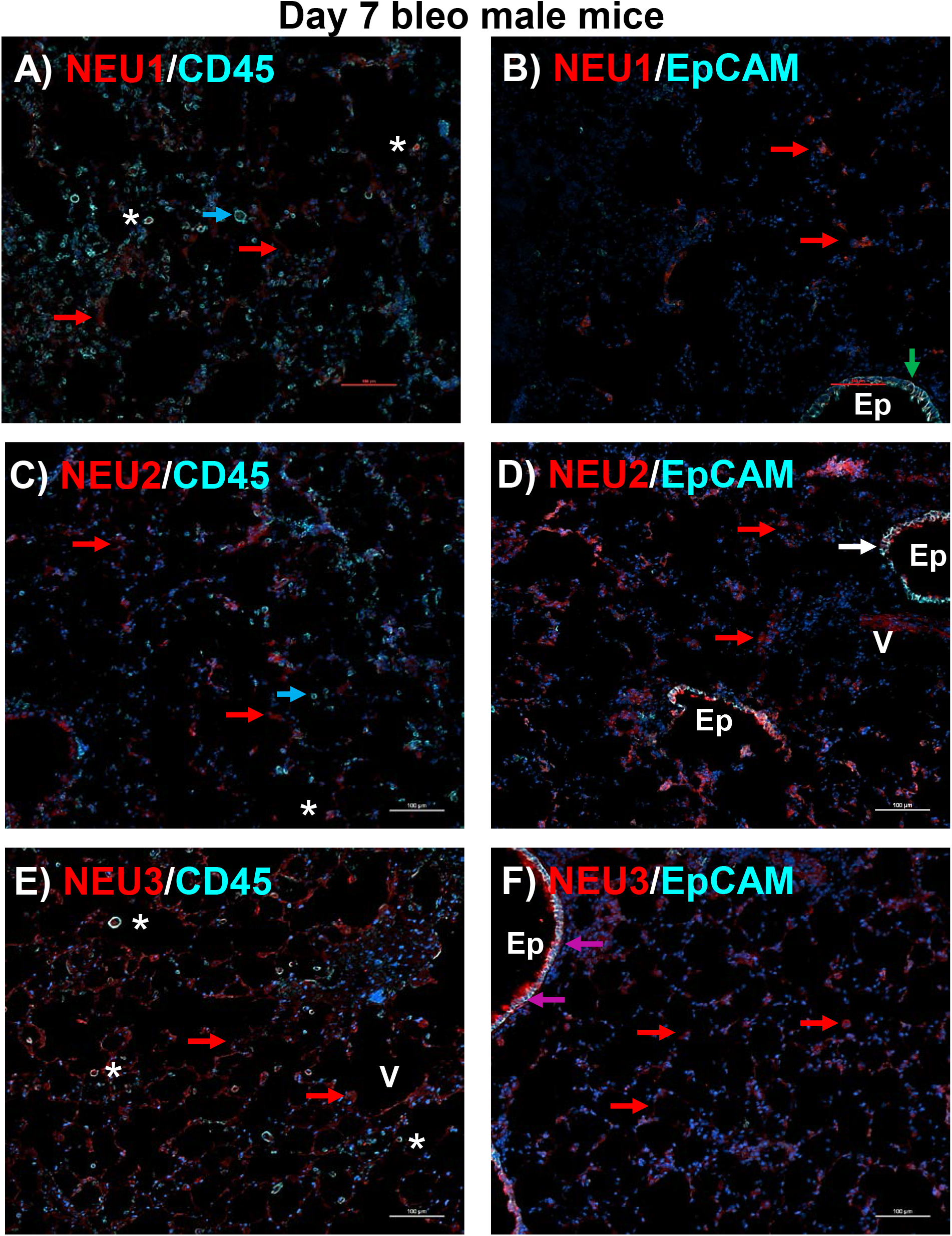
Localization of sialidases with CD45 positive immune and EpCAM positive epithelial cells in male mice at day 7. Cryosections of bleomycin treated male mouse lungs post BAL at day 7 were stained with (left panels) anti-CD45 (cyan) or (right panels) anti-EpCAM (cyan) antibodies, and either anti-NEU1, -NEU2, or -NEU3 antibodies (red), and DAPI (blue). All images are representative of three mice per group. Bars are 100 μm. Cyan arrow indicates CD45 positive cells, red arrow indicates sialidase positive cells, asterisks indicate CD45 and sialidase dual positive cells. Green arrow indicates EpCAM and NEU1 positive cells, white arrow indicated EpCAM and NEU2 positive cells, and magenta arrows indicate EpCAM and NEU3 positive cells. **Ep** indicates EpCAM positive epithelial vessels and **V** indicates EpCAM negative endothelial vessel.

**Figure 5:**
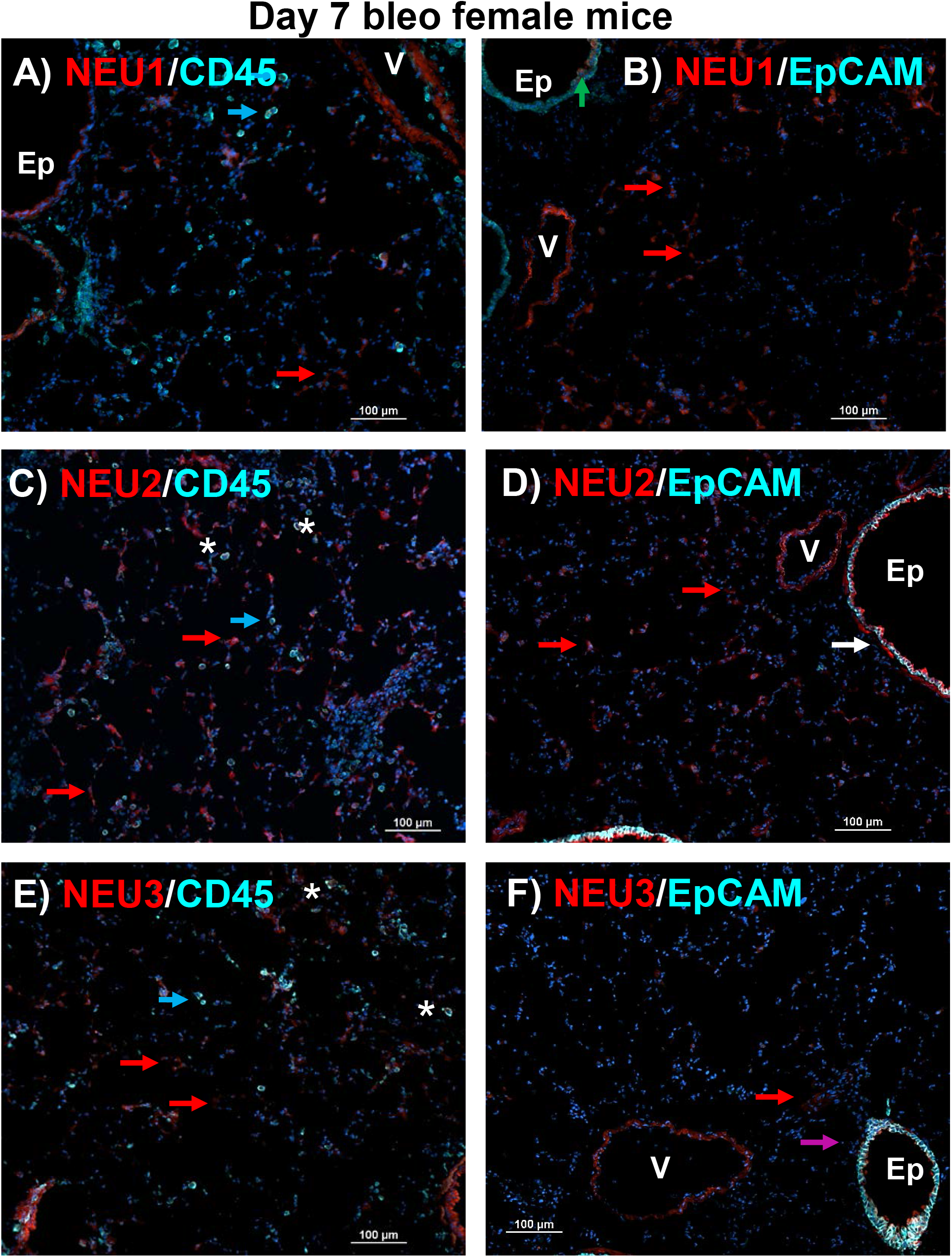
Localization of sialidases with CD45 positive immune and EpCAM positive epithelial cells in female mice at day 7. Cryosections of bleomycin treated female mouse lungs post BAL at day 7 were stained with (left panels) anti-CD45 or (right panels) anti-EpCAM (cyan) antibodies, and either anti-NEU1, -NEU2, or -NEU3 antibodies (red), and DAPI (blue). All images are representative of three mice per group. Bars are 100 μm. Cyan arrow indicates CD45 positive cells, red arrow indicates sialidase positive cells, and asterisks indicate CD45 and sialidase dual positive cells. Green arrow indicates EpCAM and NEU1 positive cells, white arrow indicates EpCAM and NEU2 positive cells, and magenta arrow indicates EpCAM and NEU3 positive cells. **Ep** indicates EpCAM positive epithelial vessels and **V** indicates EpCAM negative endothelial vessels.

**Figure 6:**
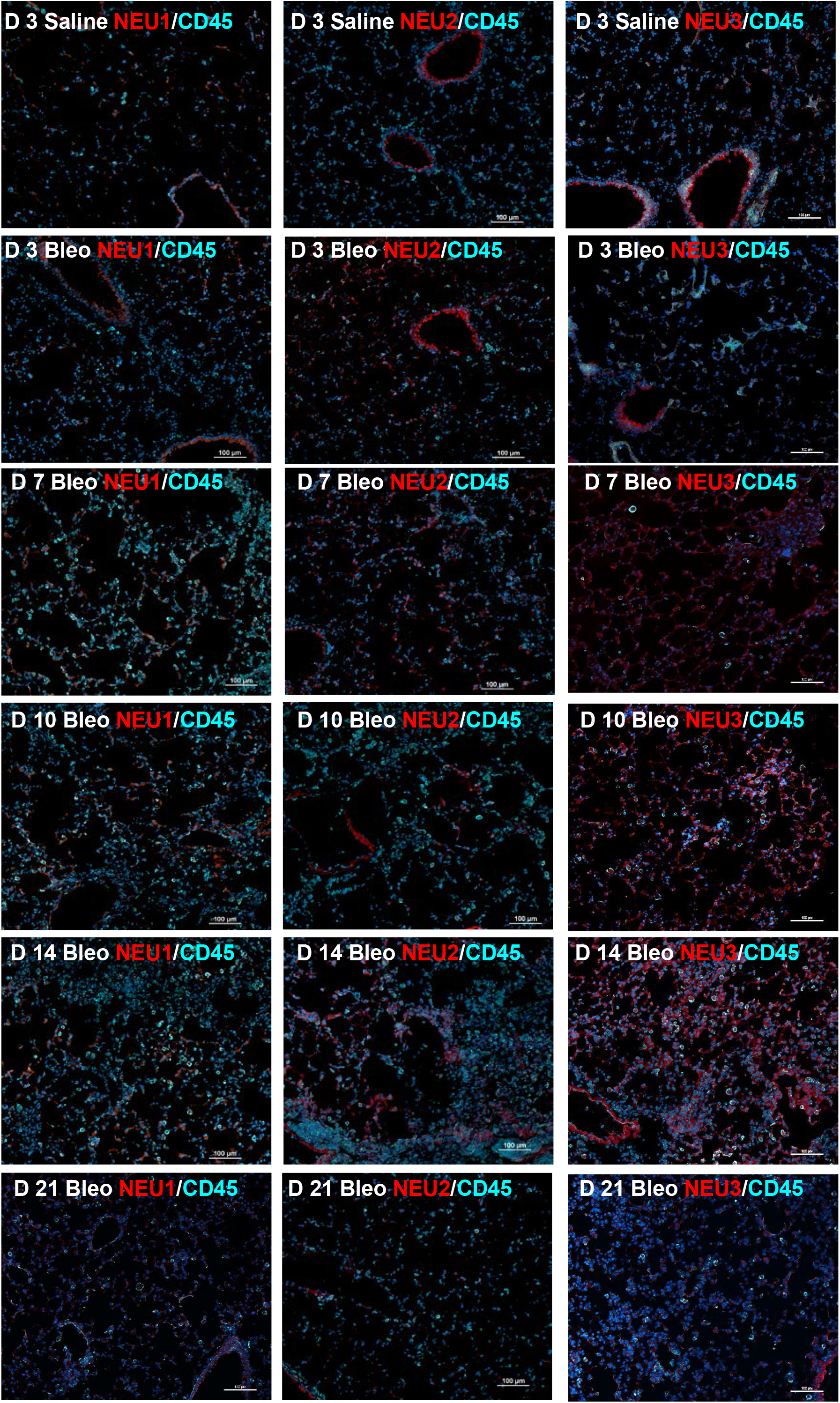
Sialidase and CD45 staining in male mice. Cryosections of saline and bleomycin treated male mouse lungs post BAL at days 3 to 21 were stained with anti-CD45 (cyan), and either anti-NEU1, -NEU2, or -NEU3 antibodies (red), and DAPI (blue). All images are representative of three mice per group. Bars are 100 μm.

In bronchiole epithelial cells of naïve mice (Figures S6) and at all time-points after saline or bleomycin treatment, NEU2 and NEU3 staining tended to be highest at the apical region of the epithelial vessels, with the EpCAM staining being more basal (white and magenta arrows and Ep labeled vessels in Figures 4 and 5 and unlabeled in Figures 6-9). We also observed NEU2 and NEU3 staining in all conditions in the EpCAM negative small endothelial vessels (marked as V in Figures 4-5), but without any apical or basal preference (Figures 4-9).

NEU2 is upregulated at day 7 in male mice (Figures 3B, 4 and S3), and these NEU2 positive cells were localized with some EpCAM positive cells (white arrows Figures 4D and 5D) and CD45 positive cells (* in Figures 4 and 5), but most of the NEU2 staining was in the alveolar cells (cells with red arrows in Figures 4 and 5, and unmarked in 6 and 8). In female mice, NEU2 staining was also mostly localized to EpCAM and CD45 negative cells (Figures 5, 7 and 9).

**Figure 7:**
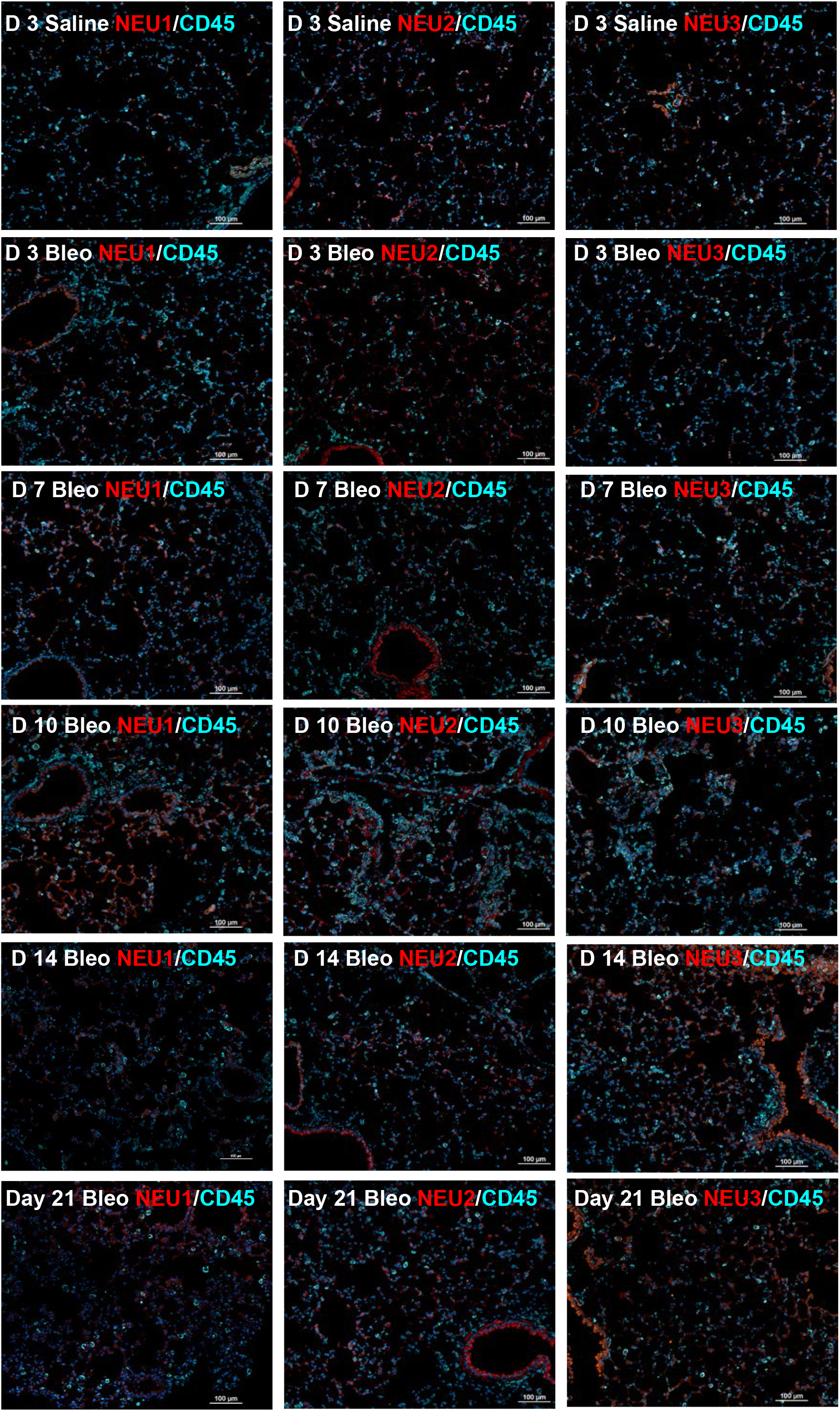
Sialidase and CD45 staining in female mice. Cryosections of saline and bleomycin treated female mouse lungs post BAL at days 3 to 21 were stained with anti-CD45 (cyan), and either anti-NEU1, -NEU2, or -NEU3 antibodies (red), and DAPI (blue). All images are representative of three mice per group. Bars are 100 μm.

**Figure 8:**
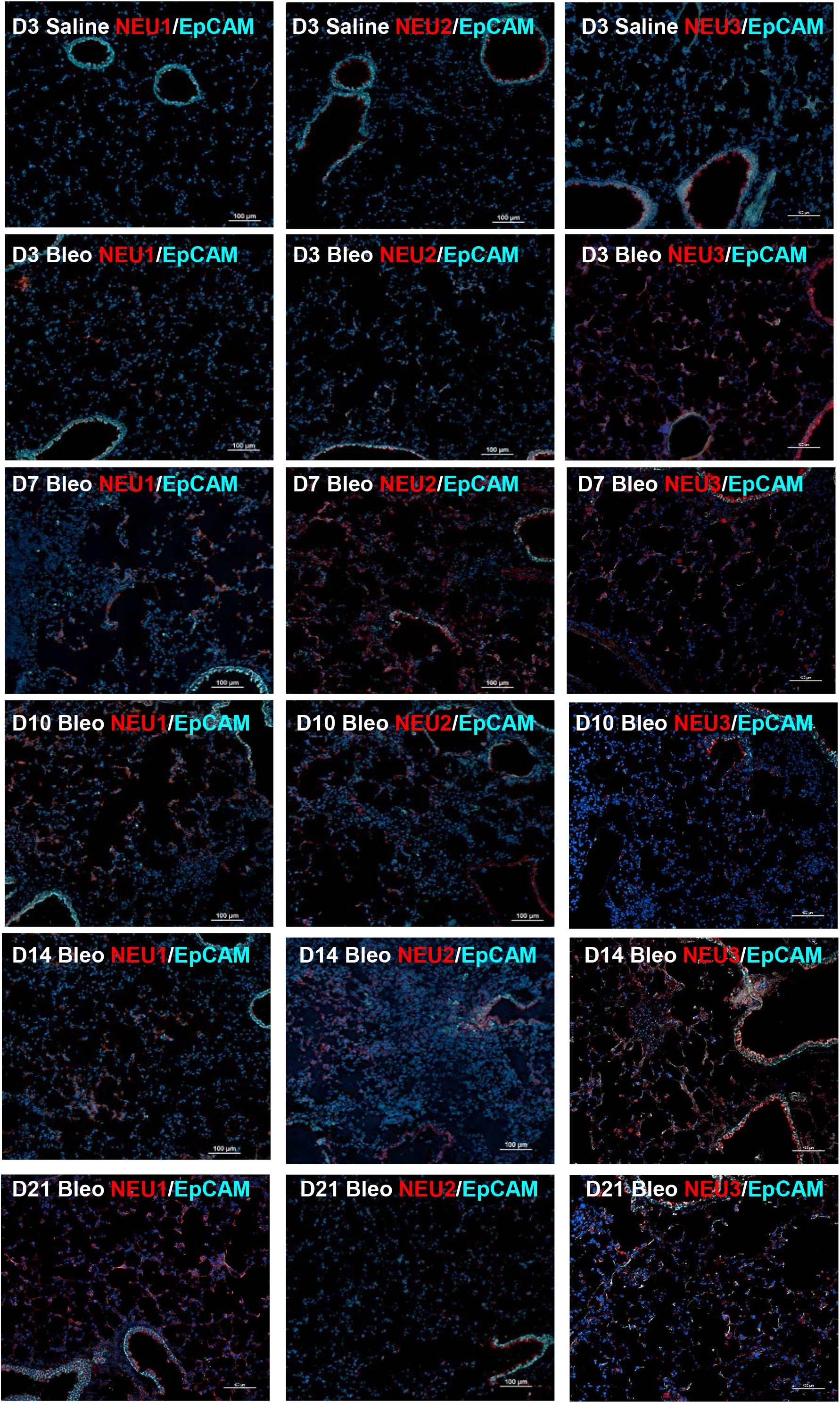
Sialidase and EpCAM staining in male mice. Cryosections of saline and bleomycin treated male mouse lungs post BAL at days 3 to 21 were stained with anti-EpCAM (cyan), and either anti-NEU1, -NEU2, or -NEU3 antibodies (red), and DAPI (blue). All images are representative of three mice per group. Bars are 100 μm.

**Figure 9:**
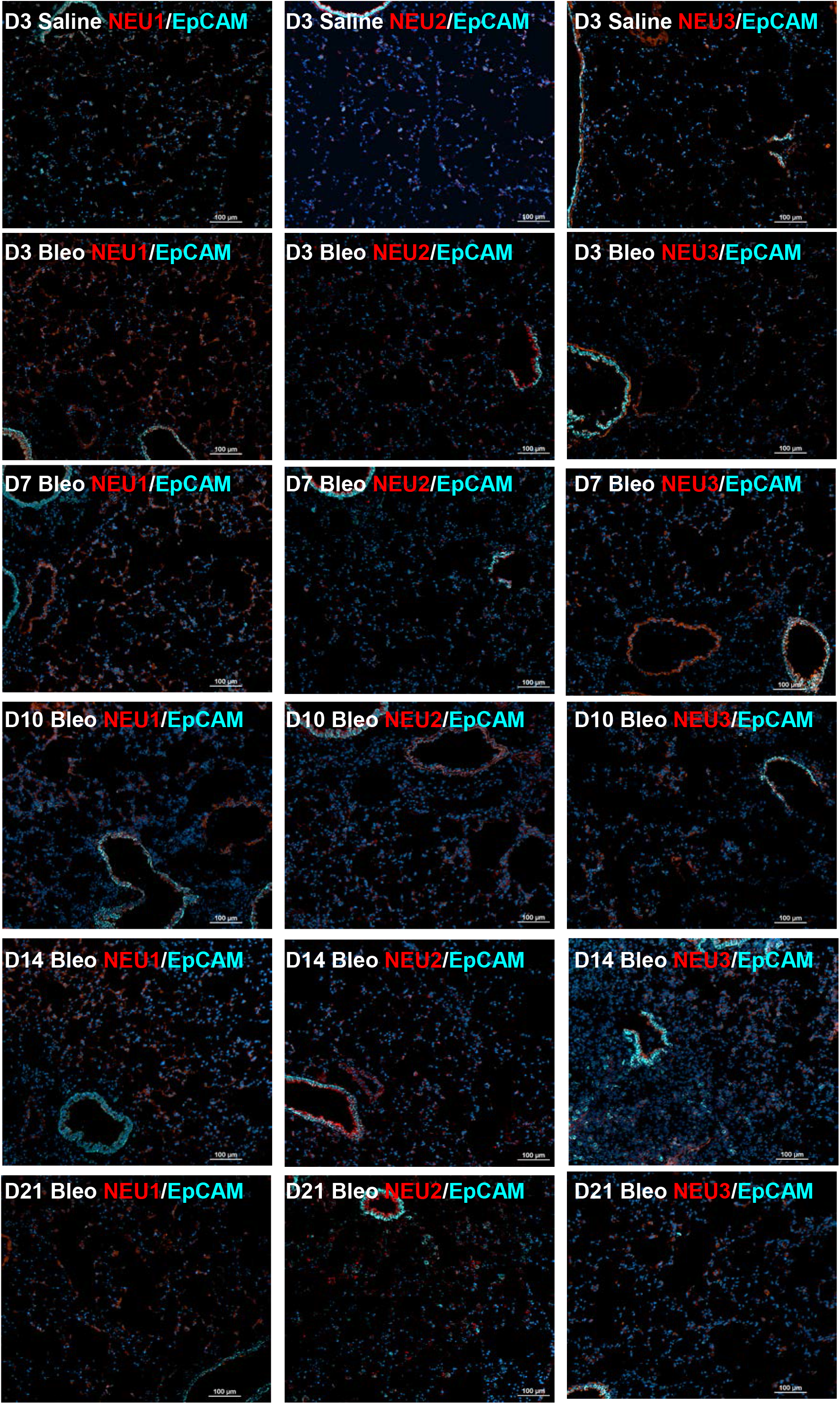
Sialidase and EpCAM staining in female mice. Cryosections of saline and bleomycin treated female mouse lungs post BAL at days 3 to 21 were stained with anti-EpCAM (cyan), and either anti-NEU1, -NEU2, or -NEU3 antibodies (red), and DAPI (blue). All images are representative of three mice per group. Bars are 100 μm.

NEU3 is generally upregulated at all time after bleomycin aspiration, and these NEU3 positive cells localized to EpCAM positive cells (magenta arrows and Ep labeled vessels in Figures 4F and 5F) and CD45 positive cells (* in Figures 4 and 5), as well as alveolar cells, in both males (cells with red arrows in Figure 4, and unmarked in 6 and 8) and females (Figure 5, and unmarked in 7 and 9). Together, these results suggest that NEU1, NEU2, and NEU3 are expressed by many cell types in the lung, both in naïve mice and after aspiration of saline or bleomycin.

## Discussion

In this report, we observed that following bleomycin aspiration, sialidase upregulation was different between male and female mice. NEU1 was upregulated at day 14 and 21 in male and day 10 in female mice. NEU2 was upregulated at day 7 in male mice and day 10 in female mice. NEU3 was elevated at days 7 and then again at days 14 and 21 in male mice but was upregulated at all time points after bleomycin in female mice. As discussed below, we observed little expression of NEU4 in male or female mouse lungs before and after bleomycin treatment. The sex-based differences in sialidase expression following bleomycin reinforce the idea that sex has a significant effect on the response of mice to bleomycin. ^18, 40-43^

Whether the changes in sialidase protein levels after bleomycin treatment are associated with changes in levels of the mRNAs encoding the sialidases is unclear. NEU1 and NEU3 protein have previously been detected in in human and mouse lung tissue by immunocytochemistry.^11, 12, 14, 45^ However, in the 14 day bleomycin model using female mice, there were increased levels of *NEU1* and *NEU2* mRNA, *NEU3* mRNA levels were unchanged, and *NEU4* mRNA was undetectable in lung tissue after bleomycin aspiration, but only NEU1 protein could be detected by western blotting.^17^ We previously observed that the profibrotic cytokine TGF-β1 increases NEU3 protein levels by decreasing NEU3 degradation and by increasing the translation of *NEU3* mRNA in polysomes, without changes in the levels of *NEU3* mRNA.^47^ Human neutrophils contain sialidase enzymic activity but express only NEU2 protein, even though *NEU1* and *NEU4* are the predominant sialidase mRNAs in neutrophils. ^48^ Sialidase protein production, secretion, and activity can also be regulated by proteins that bind to sialidases, especially protective protein/cathepsin A (PPCA) binding to NEU1.^49^ These data suggest that as sialidase mRNA levels are regulated by post-transcriptional mechanisms, and as sialidase proteins can bind chaperones such as PPCA, mRNA abundance may not be a good readout for sialidase protein expression.

The increased expression of NEU1 in day 3 saline treated male mice, compared to naïve male mice, suggests that either aspiration of saline, or the process of isoflurane anesthesia itself, was enough of a signal to induce NEU1 expression in male mice. Inhaled saline and isoflurane have both been shown to induce or modulate both lung injury and inflammatory response.^50-54^ Compared to female mice, male mice have increased airway inflammation in response to bleomycin, ^40-43^ and increased airway inflammation and decreased airway function in response to inflammatory agents such as LPS, and bronchoconstrictive agents such as methacholine.^40, 55^ These reports and our studies suggest that in male but not female mice, a very slight insult elevates levels of NEU1, and that NEU1 may mediate some aspects of inflammation.

NEU1, NEU2, and NEU3 were detectable in epithelial and endothelial cells, CD45 positive immune cells, and alveolar cells. When any of these sialidases were elevated after bleomycin, there were elevated levels in all of the above cell types, indicating that the mechanisms that elevate sialidases in response to bleomycin are not cell-type specific. The expression of sialidases in the epithelial vessels had a clear apical to basal expression pattern, with the sialidases being expressed in the apical areas. This may suggest that the epithelial cells are either secreting the sialidases into the lumen of the lung, or are retaining the sialidases in areas of the cell closest to the lumen. Either scenario would provide the cell with elevated levels of sialidases closest to the airways of the lung as a possible defense mechanism to pathogens, or to modulate the surfactants of the lung.^56^

Changes in glycosylation and the levels of sialidases are also found in other diseases.^6, 57, 58^ In cardiovascular disease models, both NEU1 and NEU3 are upregulated, and genetic disruption of *NEU1* or *NEU3* in mice, or treatment of mice with sialidase inhibitors, attenuate atherosclerosis, cardiac hypertrophy, or cardiac fibrosis.^59-61^ In cancer, all four sialidases can be upregulated, leading to increased cellular responses to growth factors such as epidermal growth factor and IL- 6, increased metastasis, and maintenance of cancer stem cells.^6, 57, 58^ In addition, the hypomorphic *NEU1* (Neu1^hypo^) mouse, with reduced levels of NEU1, has reduced inflammation in models of cardiac and hepatic disease, and *NEU3* knockout mice have a reduced incidence of colitis-associated colon cancer.^62-64^ These findings all suggest that elevated levels of one or more sialidases can promote inflammation, cellular proliferation, and fibrosis.

Together, our observation of different sialidases peaking at different times in male and female mice after bleomycin, combined with the observations of sialidases being involved in inflammation, fibrosis, and cancer in non-lung diseases, and our observation that NEU4 is sometimes upregulated and sometimes not upregulated in the bleomycin model, suggest that the four mammalian sialidases have specific roles in lung inflammation and fibrosis. The observations that of the four sialidases, only NEU3 is elevated in lung fibrosis in both male and female mice, that male and female mice lacking NEU3 essentially do not develop fibrosis in response to bleomycin, ^14^ that NEU3 inhibitors block fibrosis in both male and female mice,^13, 16^ and that aspiration of NEU3 induces fibrosis in male and female mice,^18^ suggest that a specific role of NEU3 is to promote fibrosis.

## Acknowledgments

We thank the LARR (Laboratory Animal Resources and Research) staff at Texas A&M University for animal care.

## Disclosure of interest

RHG is a scientific founder of Prosia Therapies, an early stage company developing NEU3 inhibitors as therapeutics for pulmonary fibrosis. Texas A&M University has published patent applications on the use of sialidase inhibitors (United States Patent Application 20190201485) to regulate fibrosis. DP, WC, and RHG are inventors on pending patent applications for the use of sialidase inhibitors as anti-inflammatory, anti-fibrotic, and/or anti-obesity compounds.

## Ethics approval and consent to participate

Animal experiments were carried out in strict accordance with the recommendations in the Guide for the Care and Use of Laboratory Animals of the National Institutes of Health. The protocol was approved by Texas A&M University Animal Use and Care Committee (IACUC 2020-0272).

## Availability of data and materials

The data that support the findings of this study are available from the corresponding authors upon reasonable request.

## Funding

This work was supported by National Institutes of Health National Heart, Lung, and Blood Institute [Grant R01 HL132919]. The funders had no role in study design, data collection and analysis, decision to publish, or preparation of the manuscript.

## Authors’ contributions

DP and RHG conceived and designed research; DP, KS, and WC performed experiments; DP, KS, WC and RHG analyzed data; DP and KS prepared figures; DP and RHG drafted the manuscript; DP, KS, WC, and RHG edited and revised the manuscript; all authors read and approved the final manuscript.

## Supplementary Figure Legends

**Figure S1:**
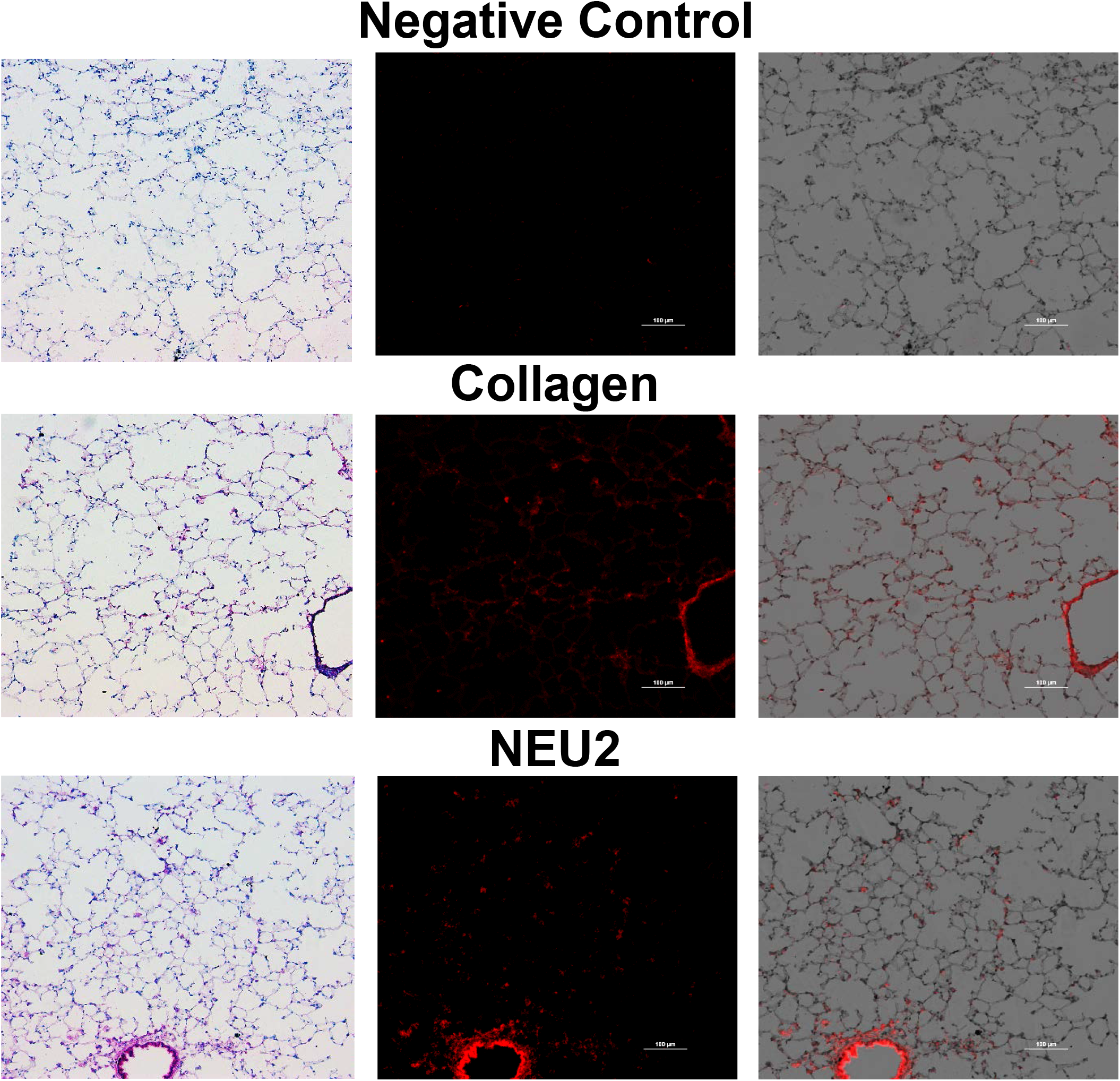
Quantification of sialidase staining by fluorescence. To quantify antibody staining developed by Vector Red Alkaline Phosphatase, bright field images (left column), fluorescent images using the Texas Red filter set (middle column), and overlap images (right column) were collected using color and monochrome cameras. Top row is negative control staining with irrelevant rabbit antibodies, middle row is positive control staining with anti-collagen-I antibodies, and bottom row is anti-NEU2 antibodies. Fluorescence intensity image quantification was done using the red fluorescent image and the total area of the tissue, from a corresponding bright field image, and these were then used to calculate the percentage area of tissue stained as a percentage of the total area of the tissue using ImageJ software. All images are representative of three mice per group. Bars are 100 μm.

**Figure S2:**
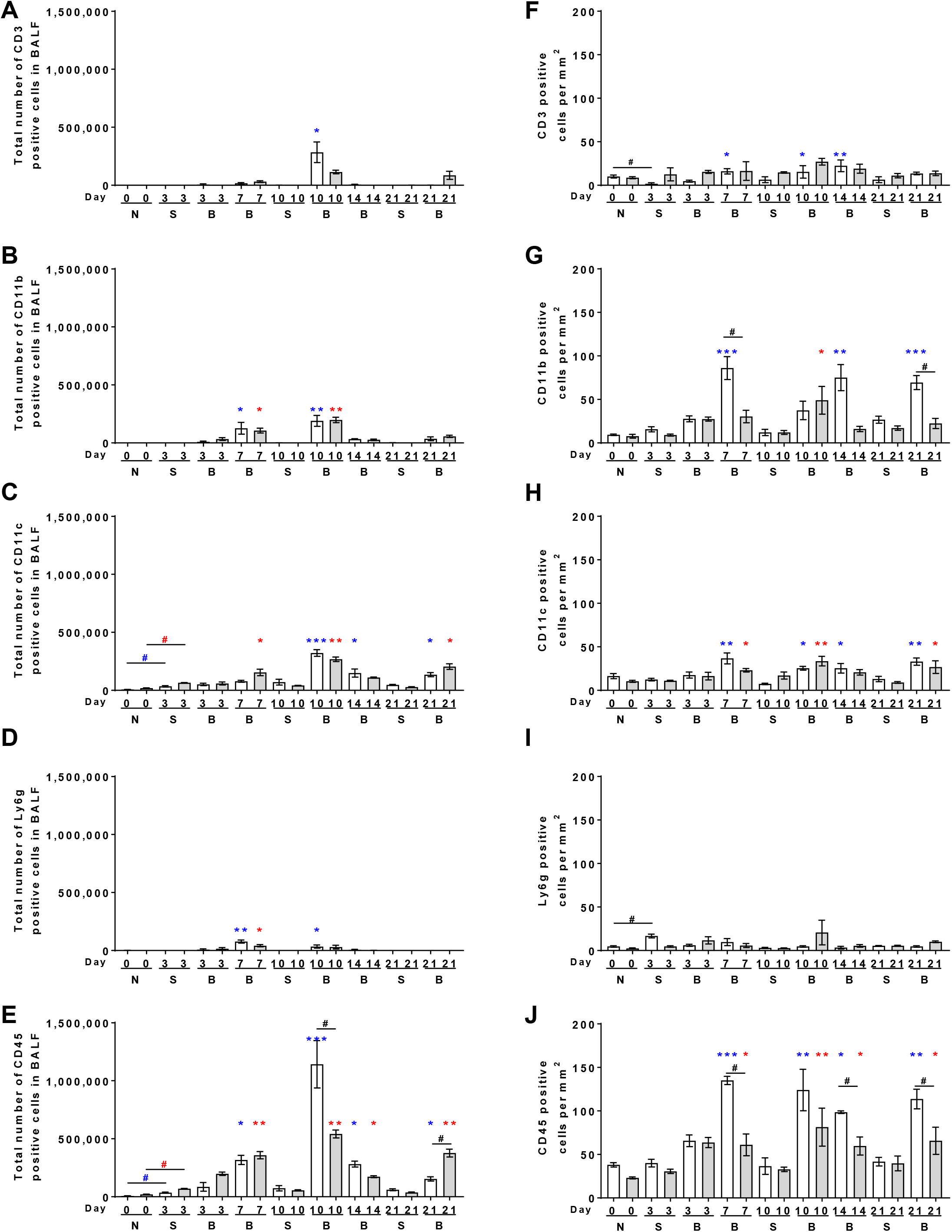
Quantification of inflammation and fibrosis in mouse lungs. Quantification of immune cells in **A-E)** BALF cell spots or **F-J)** cryosections of post-BAL lung tissue from naïve (N) untreated mice, and mice at days 3, 7, 10, 14, or 21 after saline (S) or bleomycin (B). **A-E)** BALF cell spots from male and female mice were stained for the markers CD3, CD11b, CD11c, Ly6g, and CD45. The percentage of cells stained was determined in 5 randomly chosen fields of 100–150 cells, and the percentage was multiplied by the total number of BAL cells for that mouse to obtain the total number of BAL cells staining for the marker. **F-J)** Cryosections of male and female mouse lungs were stained for CD3, CD11b, CD11c, Ly6g, and CD45, and counts were converted to the number of positive cells per mm^2^. Values are mean ± SEM, n=3 male or 3 female mice. Blue asterisks indicate significance for male mice and red asterisks indicate significance for female mice compared to day 3 saline treated mice. *p < 0.05; **p < 0.01, ***p < 0.001 (one-way ANOVA, Dunnett’s test). # p < 0.05 (t test).

**Figure S3:**
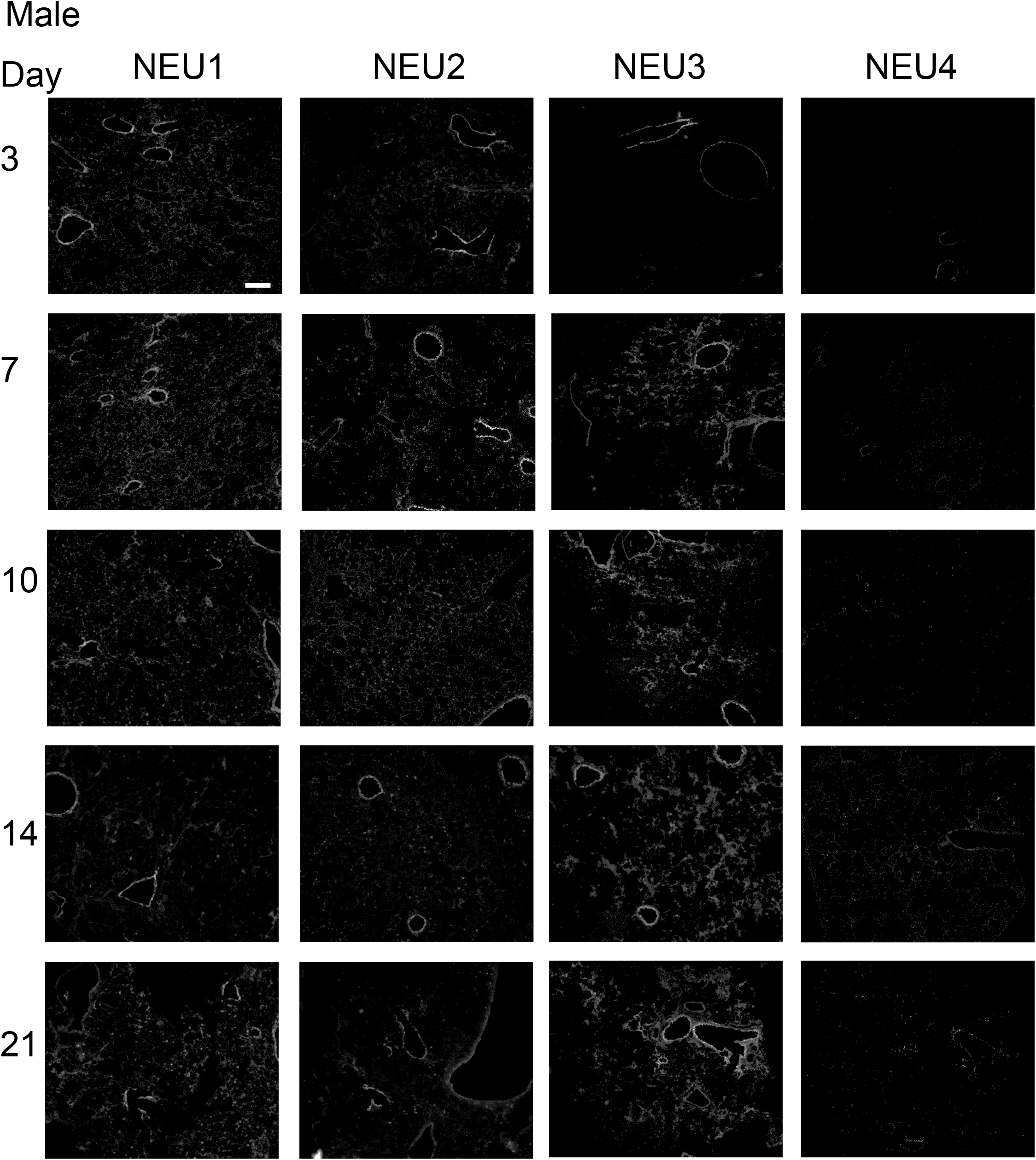
Vector red fluorescent sialidase staining in male mice. Cryosections of bleomycin treated male mouse lungs post BAL at days 3 to 21 were stained with anti-sialidase antibodies, and imaged with the Texas Red filter set. All images are representative of three mice per group. Bar is 200 μm.

**Figure S4:**
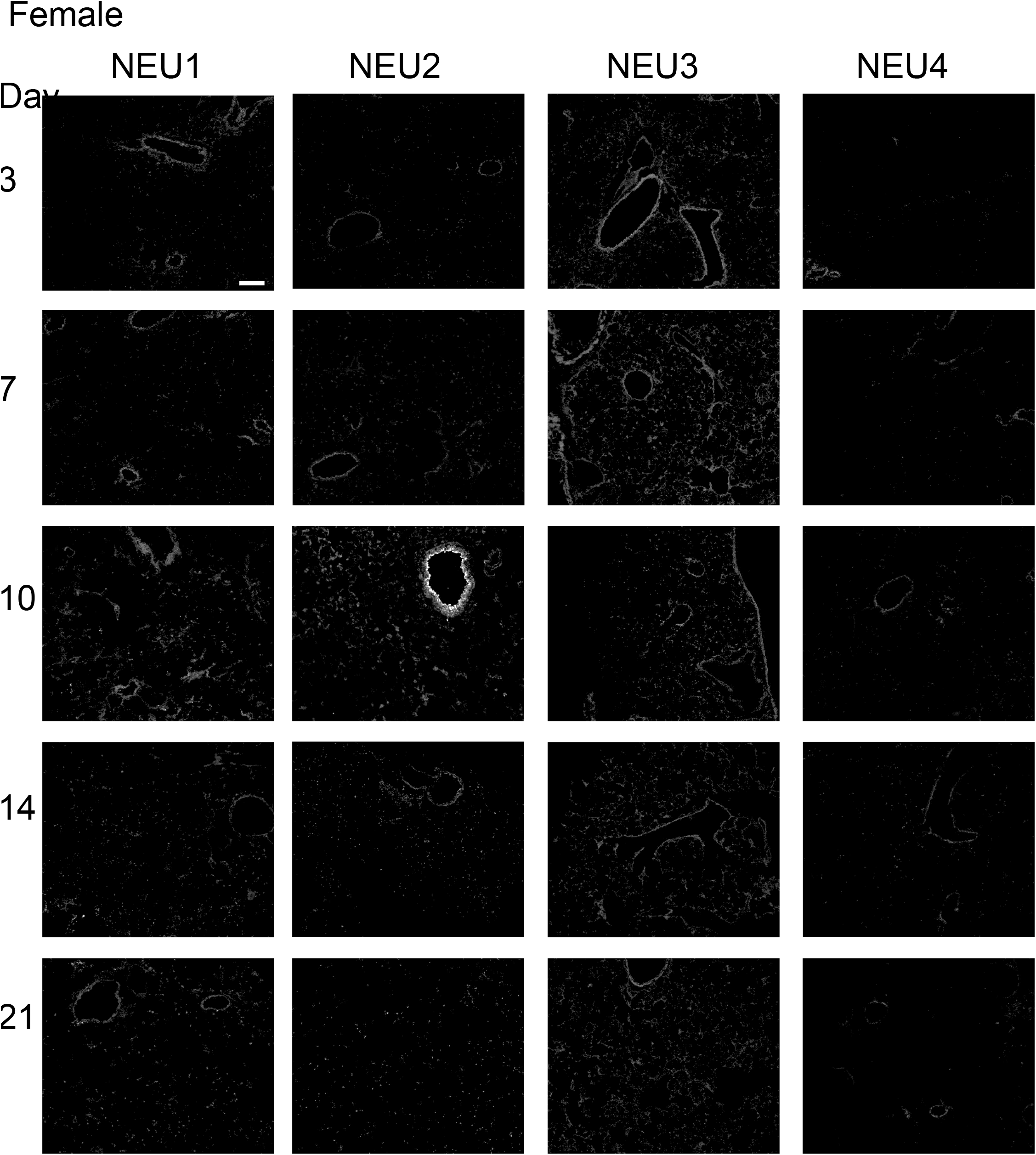
Vector red fluorescent sialidase staining in female mice. Cryosections of bleomycin treated female mouse lungs post BAL at days 3 to 21 were stained with anti-sialidase antibodies, and imaged with the Texas Red filter set. All images are representative of three mice per group. Bar is 200 μm.

**Figure S5:**
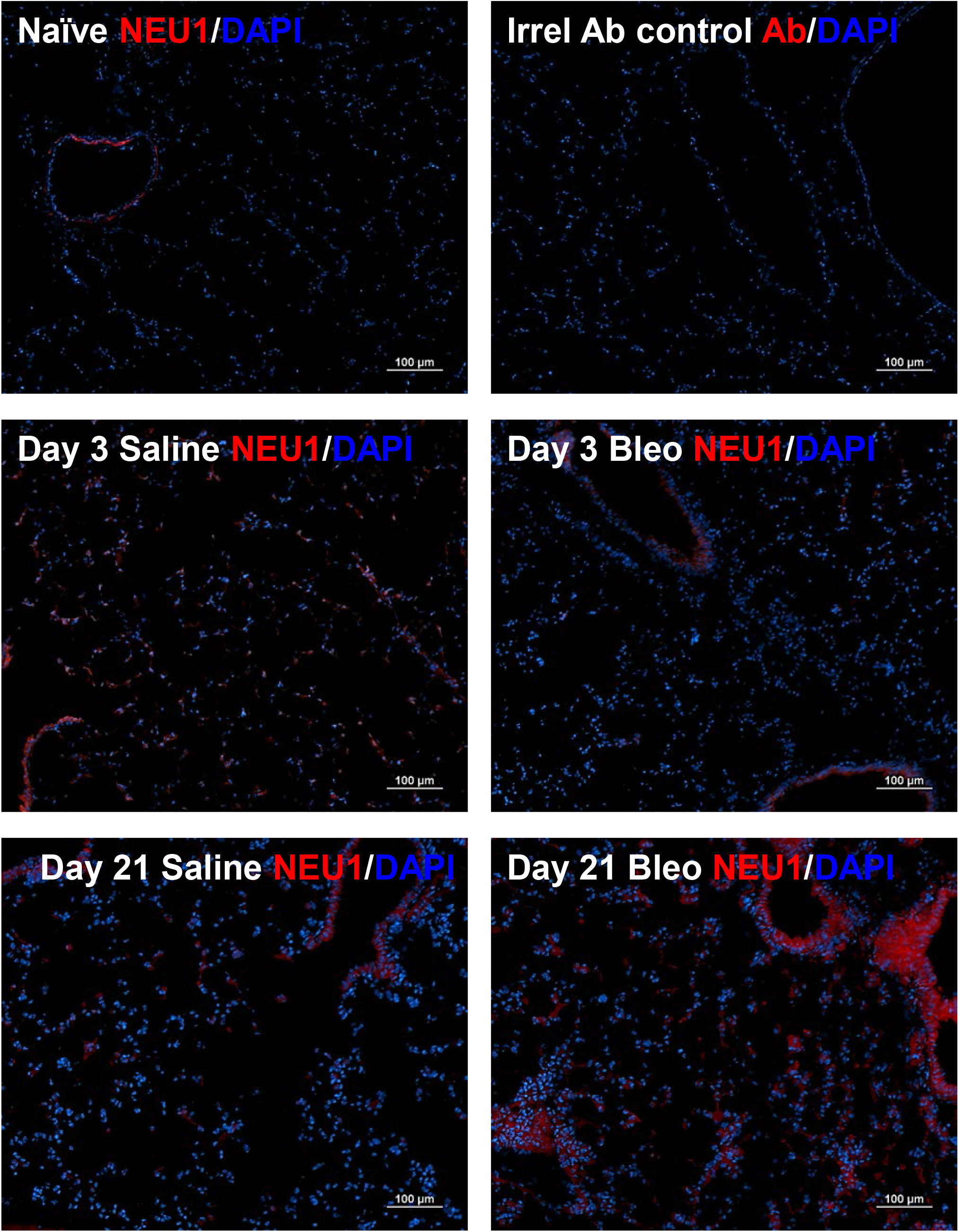
Expression of NEU1 in male mice. Cryosections of naïve, saline, and bleomycin treated male mouse lungs post BAL were stained with anti-NEU1 antibodies (red) and DAPI (blue). All images are representative of three mice per group. Bars are 100 μm.

**Figure S6:**
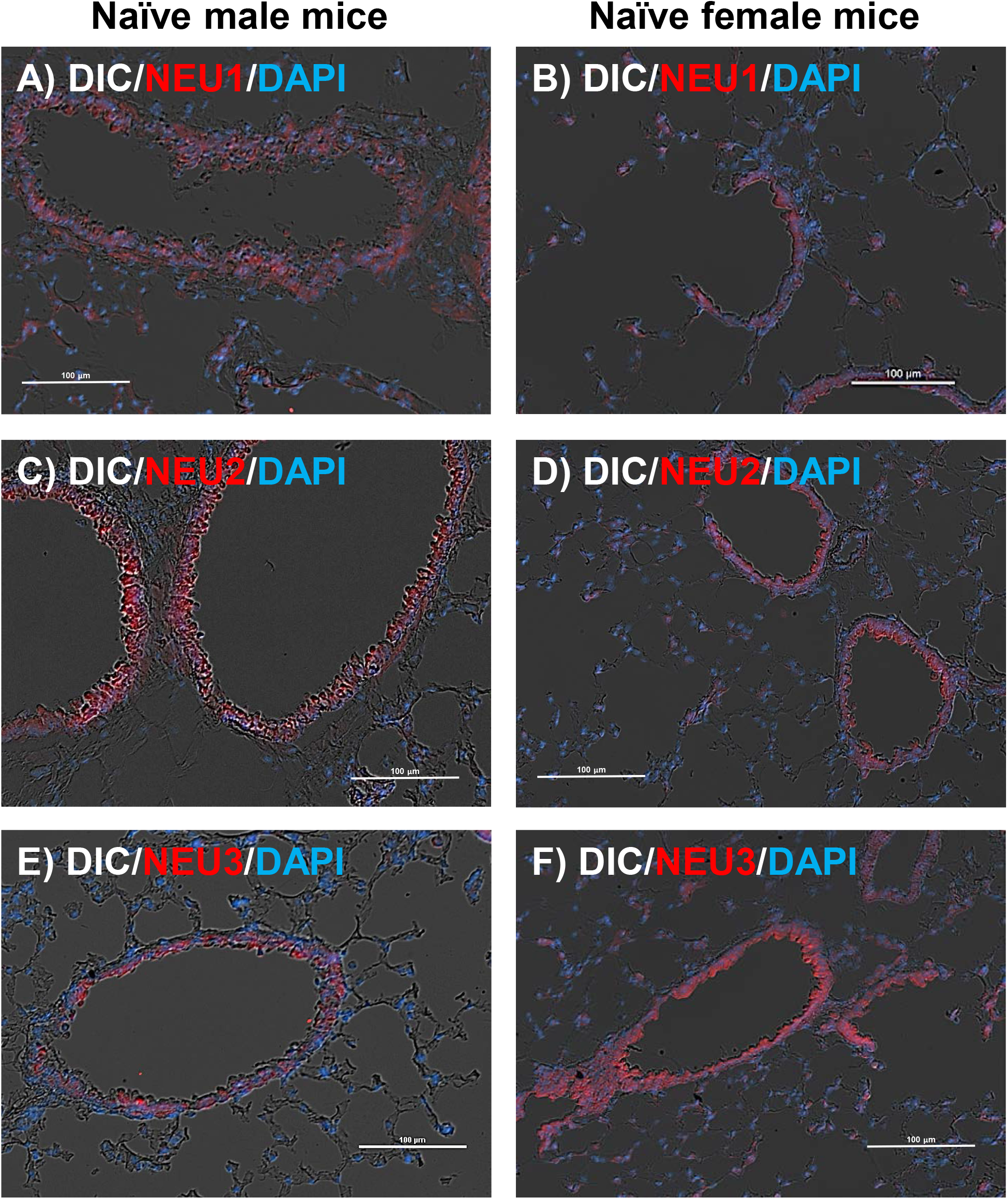
Expression of sialidases in epithelial vessels of naïve male and female mice. Cryosections of naïve **A, C, and E)** male and **B, D, and F)** female mouse lungs post BAL were stained with **A-B**) anti-NEU1 antibodies, **C-D**) anti-NEU2 antibodies, and **E-F**) anti-NEU3 antibodies. DIC and fluorescent images with sialidase antibodies (red) and DAPI (blue). All images are representative of three mice per group. Bars are 100 μm.

## References

1. Schwab I and Nimmerjahn F. Intravenous immunoglobulin therapy: how does IgG modulate the immune system? Nat Rev Immunol 2013; 13: 176–189. DOI: 10.1038/nri3401.

2. Freire-de-Lima L, Oliveira IA, Neves JL, et al. Sialic acid: a sweet swing between mammalian host and Trypanosoma cruzi. Front Immunol 2012; 3: 356. DOI: 10.3389/fimmu.2012.00356.

3. Varki A and Gagneux P. Multifarious roles of sialic acids in immunity. Ann N Y Acad Sci 2012; 1253: 16–36. DOI: 10.1111/j.1749-6632.2012.06517.x.

4. Monti E and Miyagi T. Structure and Function of Mammalian Sialidases. Top Curr Chem 2015; 366: 183–208. 2012/07/05. DOI: 10.1007/128_2012_328.

5. Smutova V, Albohy A, Pan X, et al. Structural Basis for Substrate Specificity of Mammalian Neuraminidases. PLoS ONE 2014; 9: e106320. DOI: 10.1371/journal.pone.0106320.

6. Miyagi T and Yamaguchi K. Mammalian sialidases: Physiological and pathological roles in cellular functions. Glycobiology 2012; 22: 880–896. DOI: 10.1093/glycob/cws057.

7. Zanchetti G, Colombi P, Manzoni M, et al. Sialidase NEU3 is a peripheral membrane protein localized on the cell surface and in endosomal structures. Biochemical Journal 2007; 408: 211–219. DOI: 10.1042/bj20070503.

8. Cirillo F, Ghiroldi A, Fania C, et al. NEU3 Sialidase Protein Interactors in the Plasma Membrane and in the Endosomes. J Biol Chem 2016; 291: 10615–10624. 20160317. DOI: 10.1074/jbc.M116.719518.

9. Rodriguez-Walker M and Daniotti JL. Human Sialidase Neu3 is S-Acylated and Behaves Like an Integral Membrane Protein. Sci Rep 2017; 7: 4167. 20170623. DOI: 10.1038/s41598-017-04488-w.

10. Lambré CR, Pilatte Y, Le Maho S, et al. Sialidase activity and antibodies to sialidase-treated autologous erythrocytes in bronchoalveolar lavages from patients with idiopathic pulmonary fibrosis or sarcoidosis. Clin Exp Immunol 1988; 73: 230–235.

11. Lillehoj EP, Hyun SW, Feng C, et al. Human airway epithelia express catalytically active NEU3 sialidase. Am J Physiol Lung Cell Mol Physiol 2014; 306: L876–886. 2014/03/25. DOI: 10.1152/ajplung.00322.2013.

12. Luzina IG, Lockatell V, Hyun SW, et al. Elevated expression of NEU1 sialidase in idiopathic pulmonary fibrosis provokes pulmonary collagen deposition, lymphocytosis, and fibrosis. Am J Physiol Lung Cell Mol Physiol 2016; 310: L940–954. 2016/03/20. DOI: 10.1152/ajplung.00346.2015.

13. Karhadkar TR, Pilling D, Cox N, et al. Sialidase inhibitors attenuate pulmonary fibrosis in a mouse model. Scientific Reports 2017; 7: 15069. 2017/11/10. DOI: 10.1038/s41598-017-15198-8.

14. Karhadkar TR, Chen W and Gomer RH. Attenuated pulmonary fibrosis in sialidase-3 knockout (Neu3(-/-)) mice. Am J Physiol Lung Cell Mol Physiol 2020; 318: L165–l179. 2019/10/17. DOI: 10.1152/ajplung.00275.2019.

15. Hyun SW, Liu A, Liu Z, et al. The NEU1-selective sialidase inhibitor, C9-butyl-amide-DANA, blocks sialidase activity and NEU1-mediated bioactivities in human lung in vitro and murine lung in vivo. Glycobiology 2016; 26: 834–849. 2016/05/27. DOI: 10.1093/glycob/cww060.

16. Karhadkar TR, Meek TD and Gomer RH. Inhibiting sialidase-induced TGF-β1 activation attenuates pulmonary fibrosis in mice. J Pharmacol Exp Ther 2020 2020/11/05. DOI: 10.1124/jpet.120.000258.

17. Luzina IG, Lillehoj EP, Lockatell V, et al. Therapeutic Effect of Neuraminidase-1–Selective Inhibition in Mouse Models of Bleomycin-Induced Pulmonary Inflammation and Fibrosis. Journal of Pharmacology and Experimental Therapeutics 2021; 376: 136. DOI: 10.1124/jpet.120.000223.

18. Pilling D, Sahlberg K, Karhadkar TR, et al. The sialidase NEU3 promotes pulmonary fibrosis in mice. Respiratory Research 2022; 23: 215. DOI: 10.1186/s12931-022-02146-y.

19. Cross AS, Hyun SW, Miranda-Ribera A, et al. NEU1 and NEU3 sialidase activity expressed in human lung microvascular endothelia: NEU1 restrains endothelial cell migration, whereas NEU3 does not. J Biol Chem 2012; 287: 15966–15980. DOI: 10.1074/jbc.M112.346817.

20. Moore BB and Hogaboam CM. Murine models of pulmonary fibrosis. Am J Physiol Lung Cell Mol Physiol 2008; 294: L152–160. 2007/11/13. DOI: 00313.2007[pii] 10.1152/ajplung.00313.2007.

21. Izbicki G, Segel MJ, Christensen TG, et al. Time course of bleomycin-induced lung fibrosis. International journal of experimental pathology 2002; 83: 111–119. DOI: 10.1046/j.1365-2613.2002.00220.x.

22. Tighe RM, Liang J, Liu N, et al. Recruited Exudative Macrophages Selectively Produce CXCL10 Following Non-Infectious Lung Injury. Am J Respir Cell Mol Biol 2011: 2010-0471OC. DOI: 10.1165/rcmb.2010-0471OC.

23. Misharin AV, Morales-Nebreda L, Mutlu GM, et al. Flow Cytometric Analysis of Macrophages and Dendritic Cell Subsets in the Mouse Lung. American Journal of Respiratory Cell and Molecular Biology 2013; 49: 503–510. DOI: 10.1165/rcmb.2013-0086MA.

24. Strobel B, Klein H, Leparc G, et al. Time and phenotype-dependent transcriptome analysis in AAV-TGFβ1 and Bleomycin-induced lung fibrosis models. Scientific Reports 2022; 12: 12190. DOI: 10.1038/s41598-022-16344-7.

25. Walters DM and Kleeberger SR. Mouse Models of Bleomycin-Induced Pulmonary Fibrosis. John Wiley & Sons, Inc., 2008.

26. Pilling D and Gomer RH. Persistent Lung Inflammation and Fibrosis in Serum Amyloid P Component (Apcs -/-) Knockout Mice. PLoS ONE 2014; 9: e93730. 2014/04/04. DOI: 10.1371/journal.pone.0093730.

27. Daubeuf F and Frossard N. Performing Bronchoalveolar Lavage in the Mouse. Current Protocols in Mouse Biology. John Wiley & Sons, Inc., 2012, pp.167–175.

28. Karhadkar TR, Pilling D and Gomer RH. Serum Amyloid P inhibits single stranded RNA-induced lung inflammation, lung damage, and cytokine storm in mice. PLOS ONE 2021; 16: e0245924. DOI: 10.1371/journal.pone.0245924.

29. Herlihy SE, Pilling D, Maharjan AS, et al. Dipeptidyl Peptidase IV Is a Human and Murine Neutrophil Chemorepellent. The Journal of Immunology 2013; 190: 6468–6477. 2013/05/17. DOI: 10.4049/jimmunol.1202583.

30. Pilling D, Roife D, Wang M, et al. Reduction of bleomycin-induced pulmonary fibrosis by serum amyloid P. The Journal of Immunology 2007; 179: 4035–4044. 2007/09/06. DOI: 10.4049/jimmunol.179.6.4035.

31. Haston CK, Amos CI, King TM, et al. Inheritance of susceptibility to bleomycin-induced pulmonary fibrosis in the mouse. Cancer Res 1996; 56: 2596–2601.

32. Kumar RK. Morphological methods for assessment of fibrosis. Methods MolMed 2005; 117: 179–188.

33. Hubner RH, Gitter W, El Mokhtari NE, et al. Standardized quantification of pulmonary fibrosis in histological samples. BioTechniques 2008; 44: 507-511, 514-507. 2008/05/15. DOI: 10.2144/000112729.

34. Pilling D, Zheng Z, Vakil V, et al. Fibroblasts secrete Slit2 to inhibit fibrocyte differentiation and fibrosis. Proc Natl Acad Sci U S A 2014; 111: 18291–18296. 2014/12/10. DOI: 10.1073/pnas.1417426112.

35. Pilling D, Fan T, Huang D, et al. Identification of markers that distinguish monocyte-derived fibrocytes from monocytes, macrophages, and fibroblasts. PLoS ONE 2009; 4: e7475. 2009/10/17. DOI: 10.1371/journal.pone.0007475.

36. Murdoch A, Jenkinson EJ, Johnson GD, et al. Alkaline phosphatase-fast red, a new fluorescent label. Application in double labelling for cell cycle analysis [see comments]. JImmunolMethods 1990; 132: 45–49.

37. Lauter G, Söll I and Hauptmann G. Two-color fluorescent in situ hybridization in the embryonic zebrafish brain using differential detection systems. BMC developmental biology 2011; 11: 43–43. DOI: 10.1186/1471-213X-11-43.

38. Jacobs W, Bogers J, Deelder A, et al. Adult Schistosoma mansoni worms positively modulate soluble egg antigen-induced inflammatory hepatic granuloma formation in vivo. Stereological analysis and immunophenotyping of extracellular matrix proteins, adhesion molecules, and chemokines. Am J Pathol 1997; 150: 2033–2045.

39. Schneider CA, Rasband WS and Eliceiri KW. NIH Image to ImageJ: 25 years of image analysis. Nature Methods 2012; 9: 671–675. DOI: 10.1038/nmeth.2089.

40. Voltz JW, Card JW, Carey MA, et al. Male sex hormones exacerbate lung function impairment after bleomycin-induced pulmonary fibrosis. Am J Respir Cell Mol Biol 2008; 39: 45-52. 20080214. DOI: 10.1165/rcmb.2007-0340OC.

41. Jenkins RG, Moore BB, Chambers RC, et al. An Official American Thoracic Society Workshop Report: Use of Animal Models for the Preclinical Assessment of Potential Therapies for Pulmonary Fibrosis. American journal of respiratory cell and molecular biology 2017; 56: 667–679. DOI: 10.1165/rcmb.2017-0096ST.

42. Redente EF, Jacobsen KM, Solomon JJ, et al. Age and sex dimorphisms contribute to the severity of bleomycin-induced lung injury and fibrosis. Am J Physiol Lung Cell Mol Physiol 2011; 301: L510–518. 20110708. DOI: 10.1152/ajplung.00122.2011.

43. Gilhodes J-C, Julé Y, Kreuz S, et al. Quantification of Pulmonary Fibrosis in a Bleomycin Mouse Model Using Automated Histological Image Analysis. PLOS ONE 2017; 12: e0170561. DOI: 10.1371/journal.pone.0170561.

44. Tighe RM, Liang J, Liu N, et al. Recruited Exudative Macrophages Selectively Produce CXCL10 after Noninfectious Lung Injury. American Journal of Respiratory Cell and Molecular Biology 2011; 45: 781–788. DOI: 10.1165/rcmb.2010-0471OC.

45. Lillehoj EP, Hyun SW, Feng C, et al. NEU1 sialidase expressed in human airway epithelia regulates epidermal growth factor receptor (EGFR) and MUC1 protein signaling. J Biol Chem 2012; 287: 8214–8231. DOI: 10.1074/jbc.M111.292888.

46. Ptasinski VA, Stegmayr J, Belvisi MG, et al. Targeting Alveolar Repair in Idiopathic Pulmonary Fibrosis. Am J Respir Cell Mol Biol 2021; 65: 347–365. DOI: 10.1165/rcmb.2020-0476TR.

47. Chen W, Lamb TM and Gomer RH. TGF-β1 increases sialidase 3 expression in human lung epithelial cells by decreasing its degradation and upregulating its translation. Experimental Lung Research 2020; 46: 75–80. 2020/02/28. DOI: 10.1080/01902148.2020.1733135.

48. Hyun SW, Feng C, Liu A, et al. Altered sialidase expression in human myeloid cells undergoing apoptosis and differentiation. Sci Rep 2022; 12: 14173. 20220819. DOI: 10.1038/s41598-022-18448-6.

49. Bonten EJ, Annunziata I and d’Azzo A. Lysosomal multienzyme complex: pros and cons of working together. Cellular and molecular life sciences : CMLS 2014; 71: 2017–2032. 2013/12/18. DOI: 10.1007/s00018-013-1538-3.

50. Matute-Bello G, Frevert CW and Martin TR. Animal models of acute lung injury. American Journal of Physiology - Lung Cellular and Molecular Physiology 2008; 295: L379–L399. DOI: 10.1152/ajplung.00010.2008.

51. Englert JA, Macias AA, Amador-Munoz D, et al. Isoflurane Ameliorates Acute Lung Injury by Preserving Epithelial Tight Junction Integrity. Anesthesiology 2015; 123: 377–388. DOI: 10.1097/aln.0000000000000742.

52. Chiang N, Schwab JM, Fredman G, et al. Anesthetics impact the resolution of inflammation. PLoS ONE 2008; 3: e1879.

53. Li J-t, Wang H, Li W, et al. Anesthetic Isoflurane Posttreatment Attenuates Experimental Lung Injury by Inhibiting Inflammation and Apoptosis. Mediators of Inflammation 2013; 2013: 108928. DOI: 10.1155/2013/108928.

54. Hung CJ, Liu FY, Shaiu YC, et al. Assessing transient pulmonary injury induced by volatile anesthetics by increased lung uptake of technetium-99m hexamethylpropylene amine oxime. Lung 2003; 181: 1–7. DOI: 10.1007/s00408-002-0109-4.

55. Card JW, Carey MA, Bradbury JA, et al. Gender Differences in Murine Airway Responsiveness and Lipopolysaccharide-Induced Inflammation. The Journal of Immunology 2006; 177: 621. DOI: 10.4049/jimmunol.177.1.621.

56. Hsieh I-N, De Luna X, White MR, et al. The Role and Molecular Mechanism of Action of Surfactant Protein D in Innate Host Defense Against Influenza A Virus. Frontiers in Immunology 2018; 9. Review. DOI: 10.3389/fimmu.2018.01368.

57. Glanz VY, Myasoedova VA, Grechko AV, et al. Sialidase activity in human pathologies. European Journal of Pharmacology 2019; 842: 345–350. DOI: https://doi.org/10.1016/j.ejphar.2018.11.014.

58. Miyagi T and Yamamoto K. Sialidase NEU3 and its pathological significance. Glycoconjugate Journal 2022. DOI: 10.1007/s10719-022-10067-7.

59. Demina EP, Smutova V, Pan X, et al. Neuraminidases 1 and 3 Trigger Atherosclerosis by Desialylating Low-Density Lipoproteins and Increasing Their Uptake by Macrophages. Journal of the American Heart Association 2021; 10: e018756. DOI: doi:10.1161/JAHA.120.018756.

60. Guo Z, Tuo H, Tang N, et al. Neuraminidase 1 deficiency attenuates cardiac dysfunction, oxidative stress, fibrosis, inflammatory via AMPK-SIRT3 pathway in diabetic cardiomyopathy mice. Int J Biol Sci 2022; 18: 826–840. 20220101. DOI: 10.7150/ijbs.65938.

61. Chen QQ, Ma G, Liu JF, et al. Neuraminidase 1 is a driver of experimental cardiac hypertrophy. European heart journal 2021; 42: 3770–3782. DOI: 10.1093/eurheartj/ehab347.

62. White EJ, Gyulay G, Lhoták Š, et al. Sialidase down-regulation reduces non-HDL cholesterol, inhibits leukocyte transmigration, and attenuates atherosclerosis in ApoE knockout mice. J Biol Chem 2018; 293: 14689–14706. 2018/08/12. DOI: 10.1074/jbc.RA118.004589.

63. Heimerl M, Sieve I, Ricke-Hoch M, et al. Neuraminidase-1 promotes heart failure after ischemia/reperfusion injury by affecting cardiomyocytes and invading monocytes/macrophages. Basic Res Cardiol 2020; 115: 62. 20200925. DOI: 10.1007/s00395-020-00821-z.

64. Yamaguchi K, Shiozaki K, Moriya S, et al. Reduced Susceptibility to Colitis-Associated Colon Carcinogenesis in Mice Lacking Plasma Membrane-Associated Sialidase. PLoS ONE 2012; 7: e41132. DOI: 10.1371/journal.pone.0041132.

